# Comparative transcriptomics provides insight into the evolution of cold response in Pooideae

**DOI:** 10.1101/151431

**Authors:** Lars Grønvold, Marian Schubert, Simen R. Sandve, Siri Fjellheim, Torgeir R. Hvidsten

**Author notes:** Contributed equally.

## Abstract

**Background:** Understanding how complex traits evolve through adaptive changes in gene regulation remains a major challenge in evolutionary biology. Over the last ~50 million years, Earth has experienced climate cooling and ancestrally tropical plants have adapted to expanding temperate environments. The grass subfamily Pooideae dominates the grass flora of the temperate regions, but conserved cold-response genes that might have played a role in the cold adaptation to temperate climate remain unidentified.

**Results:** To establish if molecular responses to cold are conserved throughout the Pooideae phylogeny, we assembled the transcriptomes of five species spanning early to later diverging lineages, and compared short-and long-term cold response in orthologous genes based on gene expression data. We confirmed that most genes previously identified as cold responsive in barley also responded to cold in our barley experiment. Interestingly, comparing cold response across the lineages using 8633 high confidence ortholog groups revealed that nearly half of all cold responsive genes were species specific and more closely related species did not share higher numbers of cold responsive genes than more distantly related species. Also, the previously identified cold-responsive barley genes displayed low conservation of cold response across species. Nonetheless, more genes than expected by chance shared cold response, both based on previously studied genes and based on the high confidence ortholog groups. Noticeable, all five species shared short-term cold response in nine general stress genes as well as the ability to down-regulate the photosynthetic machinery during cold temperatures.

**Conclusions:** We observed widespread lineage specific cold response in genes with conserved sequence across the Pooideae phylogeny. This is consistent with phylogenetic dating and historic temperature data which suggest that selection pressure resulting from dramatic global cooling must have acted on already diverged lineages. To what degree lineage specific evolution acted primarily through gain or loss of cold response remains unclear, however, phylogeny-wide conservation of certain genes and processes indicated that the last common ancestor may have possessed some cold response.

## Background

Adaptation to a changing climate is essential for long term evolutionary success of plant lineages. During the last ~50 million years of climate cooling, several plant species adapted to temperate regions. A key step in this transitioning was the integration of novel temperate climate cues, such as seasonal fluctuations in temperature, in the regulatory network controlling cold stress responses. Here we used the temperate grass subfamily Pooideae as a model system for studying the evolution of gene expression responses to cold stress.

The temperate grass flora is dominated by members of the subfamily Pooideae [1], and the most extreme cold environments are inhabited by Pooideae species. The ancestors of this group were, however, most likely adapted to tropical or subtropical climates [2, 3]. Many Pooideae species experience cold winters (Fig. 1ab) and although a recent study inferred adaptation to cooler environments at the base of the Pooideae phylogeny [4], it is still not known whether the Pooideae’s most recent common ancestor (MRCA) already was adapted to cold stress, or if adaptation to cold evolved independently in the Pooideae lineages.

Pooideae is a large subfamily comprising 4200 species [5], amongst them economically important species such as wheat and barley. Given the commercial importance of this group, various aspects of adaptation to temperate climate such as flowering time, cold acclimation, and frost and chilling tolerance have been studied (reviewed by [6–13]). These studies are, however, confined to a handful of species in the species rich clade “core Pooideae” [14] and recently also to its sister clade, containing the model grass *Brachypodium distachyon* [15–17]. It is thus unknown how adaptation to temperate climate evolved in earlier diverging Pooideae lineages and if conserved cold response genes could have promoted the success of this subfamily in temperate regions.

Environmental stress is assumed to be a strong evolutionary force, and the colonization of temperate biomes by Pooideae was likely accompanied by adaptation to cold conditions. A MRCA already adapted to cold (the ancestral hypothesis) offers a plausible basis for the ecological success of the Pooideae subfamily in the northern temperate regions [1]. However, paleoclimatic reconstructions infer a generally warm climate, and a very limited abundance of temperate environments, during the time of Pooideae emergence, around 50 million years ago (Mya) [18–22]. Indeed, it was not before ca. 33.5 Mya, during the Eocene-Oligocene (E-O) transition, that the global climates suddenly began to cool [23, 24] (Fig. 1c). Climate cooling at the E-O transition coincided with the emergence of many temperate plant lineages [25] and may have been an important selection pressure for improved cold tolerance in Pooideae [26, 27]. If the E-O cooling event has been the major evolutionary driving force for cold adaptation in Pooideae grasses, those findings lend support for lineage specific evolution of cold adaptation (the lineage specific hypothesis), as all major Pooideae lineages had already emerged by the time of the E-O transition [2, 28] (Fig. 1ac).

**Figure 1.**
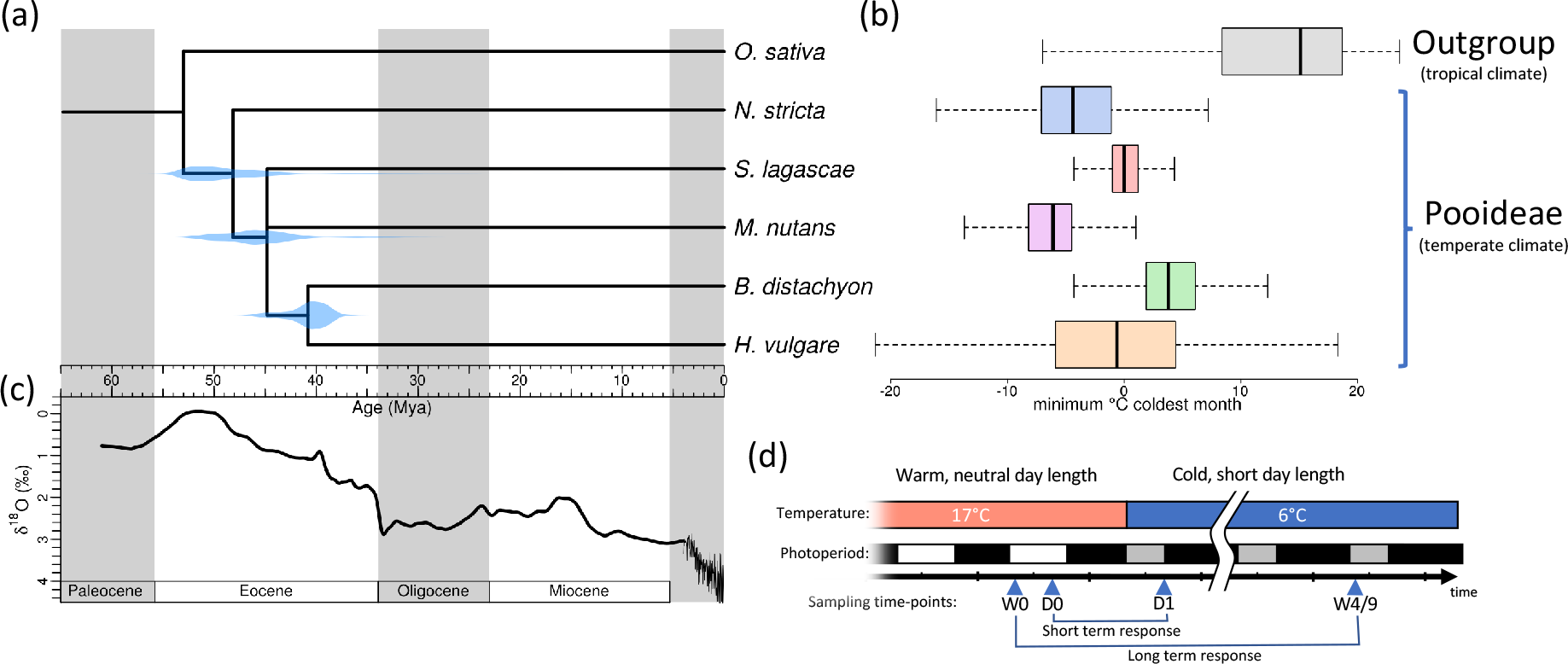
The Pooideae phylogeny, present and historic temperature data and the experimental design of this study. **(a)** Dated phylogenetic tree of the five Pooideae investigated in this study with O. sativa as an out-species. The species phylogeny was inferred from gene trees, with the distribution of mean gene-tree node ages shown in blue. (b) The range of the minimum temperature of the coldest month (WorldClim v1.4 dataset, Bioclim variable 6, 2.5 km^2^ resolution [98]) at the species geographical distribution (source: www.gbif.org). (c) Oxygen isotope ratios as a proxy for historic global temperature [18, 22] (d) Experimental design. Plants from five species of Pooideae were subjected to a drop in temperature and shorter days to induce cold response. Leaf material was sampled on the day before the onset of cold (W0 and D0), once 8 hours after cold (D1) and two times after 4 and 9 weeks (W4/W9). Short-term response was identified by contrasting gene expression in time points D0 and D1, while long-term response was identified contrasting W0 and W4/W9.

A restricted number of plant lineages successfully transitioned into the temperate region, suggesting that evolving the coordinated set of physiological changes needed to withstand low temperatures is challenging [29]. During prolonged freezing, plants need to maintain the integrity of cell membranes to avoid osmotic stress [30]. Cold and freezing tolerance is associated with the ability to cold acclimate, which is achieved through a period of extended, non-freezing cold triggered by the gradually lower temperature and day-length in the autumn. During cold acclimation, a suite of physiological changes governed by diverse molecular pathways results in an increase in the sugar content of cells, change in lipid composition of membranes and synthesis of anti-freeze proteins [13, 31]. Also, low non-freezing temperatures may affect plant cells by decreasing metabolic turnover rates, inhibiting the photosynthetic machinery and decreasing stability of biomolecules (e.g. lipid membranes) [10, 12]. Several studies have used transcriptomics to compare cold stress response, however, they focused on closely related taxa or varieties within model species [17, 32–36]. As such, these studies were not able to investigate evolutionary mechanisms underlying adaptation to cold climates of entire clades.

Here, we used *de novo* comparative transcriptomics across the Pooideae phylogeny to study the evolution of cold adaptation in Pooideae. Specifically, we aim to establish if molecular responses to cold are conserved in the Pooideae subfamily or if they are the result of lineage specific evolution. The transcriptomes of three non-model species (*Nardus stricta*, *Stipa lagascae* and *Melica nutans*), which belong to early diverging lineages, were compared to the transcriptomes of the model grass *Brachypodium distachyon* and the core Pooideae species *Hordeum vulgare* (barley). We found that only a small number of genes were cold responsive in all the investigated species, and that lineage specific evolution has been prominent in the different Pooideae lineages.

## Results

To investigate the evolution of cold response in Pooideae, we sampled leaf material in five species before and after subjecting them to a drop in temperature and shorter days (Fig. 1d). RNA-sequencing (RNA-Seq) was used to reveal the short and long term cold response of transcripts, and the conservation of these responses was analyzed in the context of ortholog groups.

### De novo transcriptome assembly identified 8633 high confidence ortholog groups

The transcriptome of each species was assembled *de novo* resulting in 146k-282k contigs, of which 68k-118k were identified as containing coding sequences (CDS, Table S1). Ortholog groups (OGs) were inferred by using the protein sequences from the five *de novo* assemblies, as well as the reference genomes of *L. perenne*, *H. vulgare*, *B. distachyon*, *Oryza sativa*, *Sorghum bicolor* and *Zea mays*. The five assembled Pooideae species were represented with at least one transcript in 24k-33k OGs (Table S1).

A set of 8633 high confidence ortholog groups (HCOGs) was identified after filtering based on gene tree topology and species representation (Table S1, Table S2). We then created a single cross species expression matrix, with HCOGs as rows and samples as columns, by summing the expression of paralogs and setting the expression of missing orthologs to zero (Table S3).

### A dated species tree of the Pooideae

A dated species tree were generated based on ortholog groups with exactly one sequence from each of the five Pooideae and rice, and using prior knowledge about the divergence times of *Oryza*-Pooideae [37] and *Brachypodium*-*Hordeum* [28] (Fig. 1a). In the most common gene tree topology, *S. lagascae* or *M. nutans* formed a monophyletic clade, but topologies where either *S. lagascae* or *M. nutans* diverged first were also common (Fig. S1). Due to this uncertainty regarding the topology, *S. lagascae* and *M. nutans* branches were collapsed to a polytomy in the consensus species tree.

### Expression clustering indicated a common global response to cold

To investigate broad scale expression patterns in cold response, we clustered all samples (including replicates) after scaling the expression values of each gene to remove differences in mean expression levels between species (Fig. 2a). This clustering revealed the differential effects of the treatments and resulted in a tree with replicates, and then time points, clustering together. Before scaling, the samples clustered by species (data not shown). An exception was time points W4 and W9, which tended to cluster together and by species, indicating that responses after 4 and 9 weeks were very similar. The fact that time points mostly clustered together before species indicated a common response to cold across species. We also observed a clear effect of the diurnal rhythm, with time points sampled in the morning (W0, W4 and W9) forming one cluster and time points sampled in the afternoon (D0 and D1) forming another. This diurnal effect might have resulted in more unreliable estimates of the long term cold response in *S. lagascae* since for this species the afternoon sample (D0) was used to replace the missing morning sample (W0).

**Figure 2.**
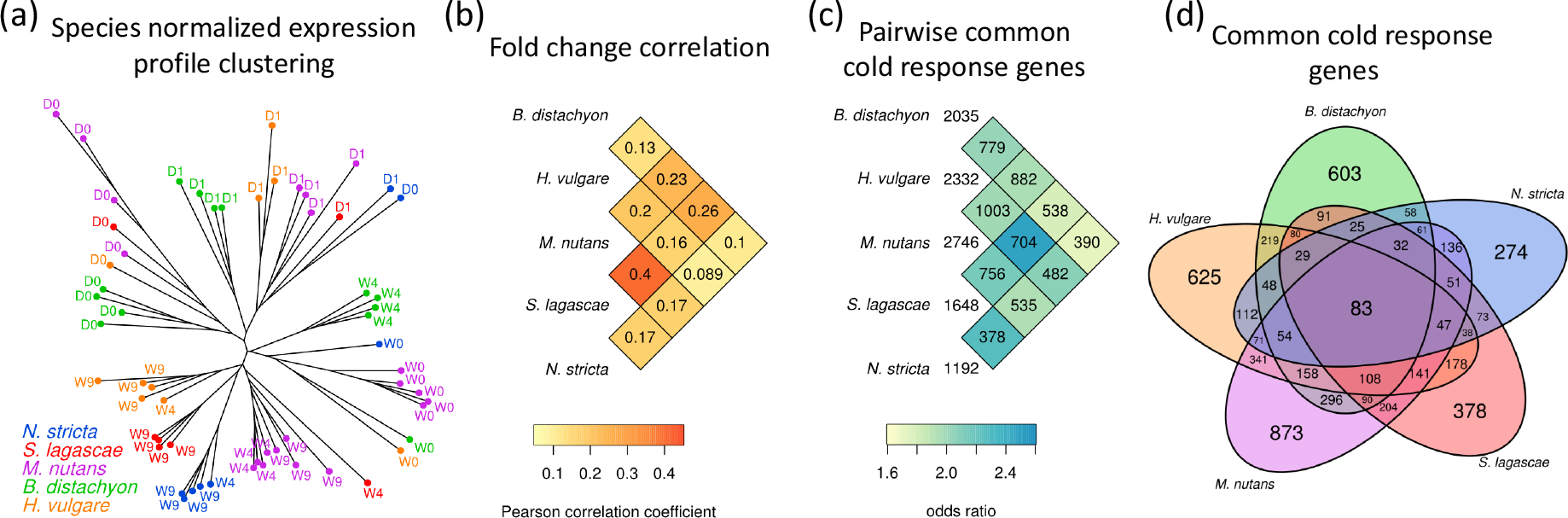
Comparison of cold response across the Pooideae. (a) Expression clustering of the samples. The tree was generated by neighbor-joining of Manhattan distances given as the sum of log fold changes between all highly expressed genes after subtracting the mean expression per species. Each tip corresponded to one sample. (b) The Pearson correlations of log fold expression changes (only short-term cold response is shown) between pairs of species. The correlations were computed based on the high confidence ortholog groups (HCOG). (c) The number of differentially expressed genes per species and shared between pairs of species. The statistical significance of the overlaps between pairs of species were indicated with odds ratios. (d) The number of differentially expressed genes in each species (FDR adjusted p-value < 0.05 and absolute fold change > 2 in either short-or long-term cold response) and overlap between species.

### Many cold responsive genes were species specific

We next examined similarities in short and long term cold response between species by analysing changes in gene expression from before cold treatment to eight hours and 4-9 weeks after cold treatment (Fig. 1d). For all species pairs, there was a low, but statistically significant, correlation between the expression fold changes of orthologs in HCOGs (Fig. 2b). A similar pattern was observed when investigating the number of orthologs classified as differentially expressed in pairs of species (FDR adjusted p-value < 0.05 and fold change > 2, Table S4, see Methods): these numbers were low compared to the number of differentially expressed genes (DEGs) in individual species, but higher than expected by chance (Fisher’s exact test p < 0.05, Fig. 2c). Finally, the number of orthologs with differential expression in more than two species were very low (Fig. 2d), with only 83 DEGs common to all five species. Noticeably, neither the similarities in differential expression nor the fold change correlations reflected the phylogenetic relationship between the species, that is, the cold responses of related species were not more similar than that of distantly related species (Fig. 2bc).

### Shared cold response genes included known abiotic stress genes

Sixteen genes shared the same cold response (short-or long-term) in the same direction (up or down) in all five Pooideae species, thus representing a response to cold that might have been conserved throughout the evolution of Pooideae (Table 1). Nine of these shared cold responsive genes belonged to families known to be involved in cold stress or other abiotic stress responses in other plant species. The most common type of response was short-term up regulation, indicating that stress response, as opposed to long-term acclimation response, is potentially more conserved.

**Table 1.** High confidence ortholog groups with conserved cold response in all five Pooideae. S = short-term response, L = long-term response. ↗ = up regulated, ↘= down regulated. Annotations were inferred from literature using orthologs. These 16 genes were the subset of the 83 genes in Fig. 2d with the same type of cold response (short-or long-term) in the same direction (up-or down-regulation) in all five species.

| Bd ortholog | Description and relation to stress response | Response |
| --- | --- | --- |
| Bradi2g39230 | **HyperOSmolality-gated CA2+ permeable channel (OSCA)** - Stress-activated calcium channels [72] that are highly conserved in eukaryotes [73]. In *Oryza sativa*, OSCA genes are differentially expressed in response to osmotic stress [74]. | $S\nearrow$ |
| Bradi2g06830 | **Calcium-binding EF-hand containing calcium exchange channel (EF-CAX)** - Calcium ions are important mediators of abiotic stress in plants [75, 76]. Expression of calcium binding proteins correlates with exposure to cold stress in several plants, e.g. *Arabidopsis thaliana* [3[30], *Musa × paradisiaca* [41] and *Hordeum vulgare* [38]. | $S\nearrow$ |
| Bradi2g05226 | **GIGANTEA** - Promotes flower development in plants [77]. In *Arabidopsis thaliana*, this gene is involved in CBF-independent freezing tolerance [78, 79], and is responsive to cold in *Zea mays* [80]. Also part of the circadian clock. | $S\nearrow$ |
| Bradi4g24967 | **Arabidopsis Pseudo-Response Regulator 3-like (AtPRR3-like)** - AtPRR3 is a member of the circadian clock quintet AtPRR1/TOC1 [81, 82]. No association to stress response found in literature. However, AtPRR3-like might be closer related to AtPRR5/9 than to AtPRR3 (See Bradi4g36077, PRR95). | $S\nearrow$ |
| Bradi2g09060 | **Triacylglycerol lipase, alpha/beta-Hydrolase superfamily** - Studies in *Arabidopsis thaliana* [83] and *Ipomoea batatas* [84] suggest that genes with alpha/beta-Hydrolase domains respond to osmotic stress. In *Triticum monococcum*, atriacylglycerol lipase was induced by pathogen stress [85]. | $S\nearrow$ |
| Bradi2g07480 | **Late-Embryogenesis-Abundant protein 14 (LEA-14)** - Responsive to drought, salt and cold stress in *Arabidopsis thaliana* [86, 87], *Betula pubescens* [88] and *Brachypodium distachyon* [89]. | $S\nearrow$ |
| Bradi1g04150 | **SNAC1-like / NAC transcription factor 67** - NAC transcription factors mediate abiotic stress responses. Osmotic stress increases the expression of SNAC1 in *Oryza sativa* [43], NAC68 in *Musa × paradisiaca* [41, 42] and NAC67 *Triticum aestivum* [44]. | $S\nearrow$ |
| Bradi4g36077 | **Pseudo-Response Regulator 95 (PRR95)** - Homologous to conserved circadian clock gene AtPRR5/9 [90, 91]. AtPRR5 gene is cold regulated in *Arabidopsis thaliana* [92] and PRR95 is cold responsive in *Zea mays* [80]. | $S\nearrow$ |
| Bradi2g43040 | **DnaJ chaperon protein** - DnaJ co-chaperons are vital in stress response and has been found to be involved in maintenance of photosystem II under chilling stress and enhances drought tolerance in tomato [93, 94]. | $S+L\nearrow$ |
| Bradi3g33080 | **Glycogenin GlucuronosylTransferase (GGT)** - GGT belongs to the GT8 protein family [95]. In *Oryza sativa*, OsGGT transcripts are induced in submerged plants and respond to various abiotic stresses except cold [96, 97]. | $L\nearrow$ |
| Bradi1g04500 | **Major facilitator superfamily transporter** - Association to stress response unknown. | $L\nearrow$ |
| Bradi3g14080 | **Glycosyl transferase** - Association to stress response unknown. | $L\nearrow$ |
| Bradi1g35357 | **Uncharacterized membrane protein** - Association to stress response unknown. | $S\nearrow$ |
| Bradi2g48850 | **Uncharacterized protein** - Association to stress response unknown. | $S\nearrow$ |
| Bradi1g33690 | **Uncharacterized protein** - Association to stress response unknown. | $S\nearrow$ |
| Bradi1g07120 | **Putative S-adenosyl-L-methionine-dependent methyltransferase** - Association to stress response unknown. | $L\searrow$ |

### Identified cold response genes confirmed previous findings

We compared the cold response genes from our data to a compilation of *H. vulgare* genes shown to be responsive to low temperature in several previous microarray studies, subsequently referred to as the Greenup genes (table S10 in [38]). We could map 33 of these 55 genes to unique OGs, of which 11 were HCOGs. We observed significant similarity in cold response between the 33 Greenup genes and the short-term cold response observed in our data (Fig. 3); for all five species (p < 0.05). However, this similarity was noticeably larger in *H. vulgare* than in the other four species. This comparison showed that our transcriptome data was consistent with previous findings in *H. vulgare*, and that cold response genes identified in *H. vulgare* exhibits some cold response in other Pooideae.

**Figure 3.**
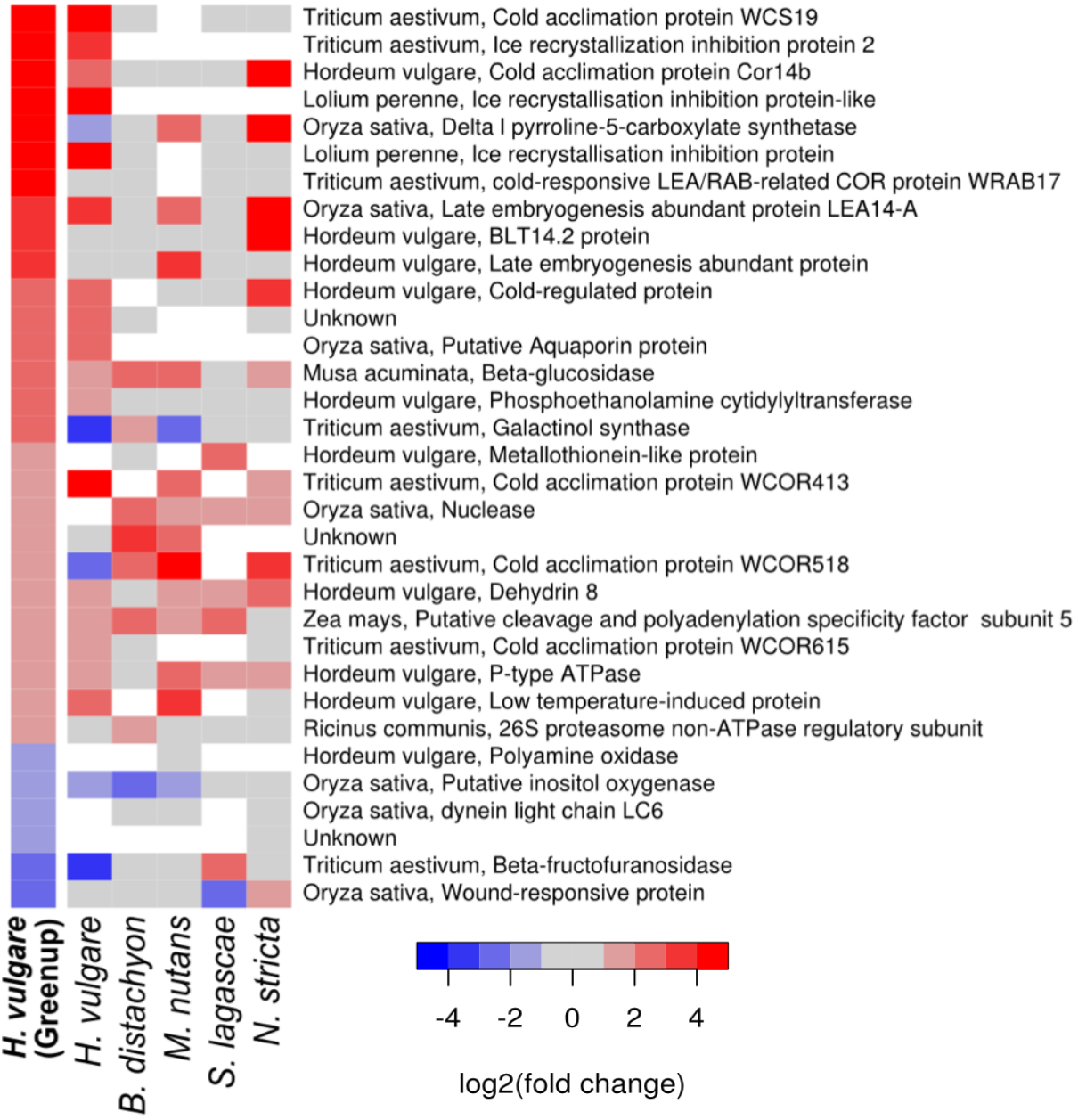
Comparison of cold response to previous studies. A reference set of H. vulgare genes independently shown to respond to cold in several studies [38] is compared to our data using short-term log fold change values. White cells represent missing orthologs. Grey cells represent orthologs that were not differentially expressed (not DEGs).

### Photosynthesis was down-regulated under cold stress

To identify biological processes that evolved regulation during different stages of Pooideae evolution, we targeted gene sets that were exclusively differentially expressed in all species within a clade in the phylogentic tree (i.e. branch specific DEGs), and tested these for enrichment of Gene Ontology (GO) biological process annotations (Fig. 4a). For the genes that were differentially expressed in all our species (Pooideae base [PB] in Fig 4b), we found enrichments for annotations related to response to abiotic stimulus, photosynthesis and metabolism. Dividing the branch specific DEGs into up- or down-regulated genes revealed up-regulation of signal transduction (two pseudo response regulators and diacylglycerol kinase 2 (DGK2)) and abiotic stimulus (Gigantea, LEA-14, DnaJ and DGK2), and down-regulation of photosynthesis and metabolism. For the genes that were exclusively differentially expressed in all species except *N. stricta* (early split [ES] in Fig. 4b), down-regulated genes were again enriched for GO annotations related to metabolism and photosynthesis.

**Figure 4.**
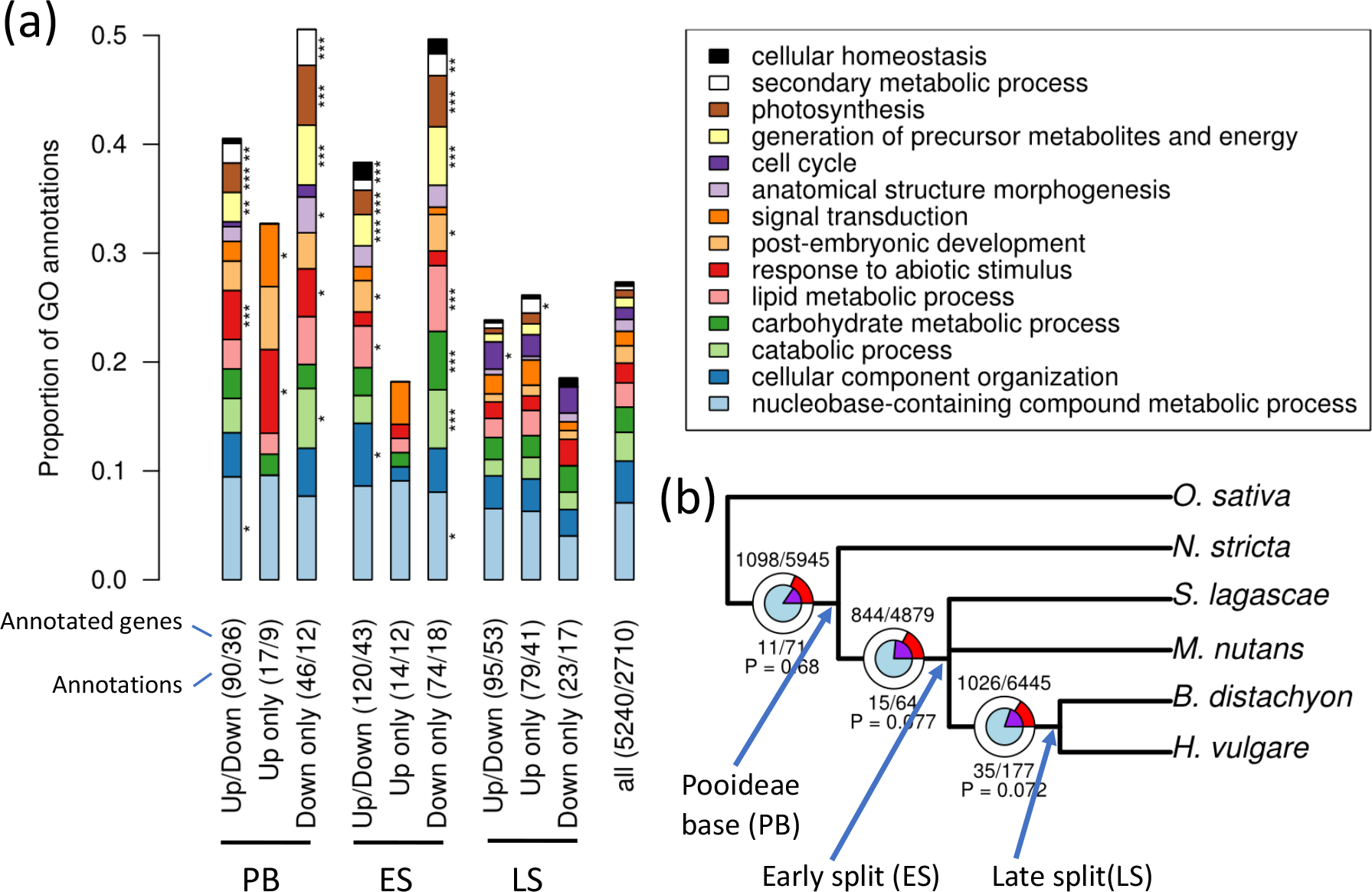
Gene Ontology enrichment and positive selection in branch specific cold responsive genes. (a) Gene ontology enrichment analysis of high confidence ortholog groups (HCOG) that were differentially expressed (DEGs) in all species (PB), only in species after N. stricta split off (ES) or only in B. distachyon and H. vulgare (LS) (* P < 0.05, ** P < 0.01, *** P < 0.005, Fisher’s exact test). Both the number of annotated genes and the number of annotations were indicated for each set of branch specific DEGs. (b) Positive selection at different stages in Pooideae evolution. The circles represent the high confidence ortholog groups that were tested for positive selection at each split (see Methods for the criteria). The inner blue circle represented HCOGs with branch specific differential expression (i.e. with genes that were cold responsive exclusively in the species under the respective branch) while the outer circle represented all other HCOGs. The purple and red pie-slices represented the proportions of HCOGs with positive selection (P < 0.05). The P-value indicated the overrepresentation of positive selection among the branch specific DEGs (Hypergeometric test).

### Cold response genes were associated with positive selection on amino acid content

For each HCOG, we tested for positive selection in coding sequences at each of the internal branches of the species tree. The tests were only performed on the branches where the gene tree topology was compatible with the species tree topology. 16-18% of the HCOGs showed significant signs of positive selection (P < 0.05) depending on the branch (Fig. 4b). Next, we tested for overrepresentation of positive selection among the branch specific DEGs. There was a tendency that gain of cold response was associated with positive selection at the early split (ES) and late split (LS) branches (P = 0.077 and P = 0.072, respectively) (Fig. 4b).

## Discussion

The ecological success of the Pooideae subfamily in the northern temperate regions must have critically relied on adaptation to colder temperatures. However, it is unclear how this adaptation evolved within Pooideae. To test whether molecular responses to cold are conserved in the Pooideae subfamily, we applied RNA-seq to identify short-and long-term cold responsive genes in five Pooideae species ranging from early diverging lineages to core Pooideae species. Since three of the species lacked reference genomes, we employed a *de novo* assembly pipeline to reconstruct the transcriptomes. We showed that this pipeline could recover a set of *H. vulgare* genes previously identified as cold responsive and that most of these genes were also cold responsive in our data (Fig. 3). In order to compare the five transcriptomes, we compiled a set of 8633 high confidence ortholog groups (HCOGs) with resolved gene tree topologies. Clustering of scaled expression data based on these ortholog groups arranged samples according to replicates, then time points and finally species, indicating that cold response represents a distinct signal in the data and confirming the soundness of the approach (Fig. 2a).

### Lineage specific adaptations to cold climates

Based on conserved genes in the 8,633 HCOGs, substantial portions of the individual Pooideae transcriptomes responded to cold (1000-3000 genes, ~10-30%). Although pairs of species shared a statistically significant number of cold responsive genes, nearly half of all responsive genes were species specific, with closely related species sharing approximately the same fraction of cold responsive genes as more distantly related species (no phylogenetic pattern, Fig. 2c-d). Moreover, only a small number of genes responded to cold in all the investigated species (83 genes, Fig. 2d). Even fewer genes responded similarly to cold in all species (e.g. short-term up-regulation, 16 genes, Table 1) and these shared cold response genes primarily included general abiotic stress genes clearly not representative of all the different molecular pathways constituting a fully operational cold response program. We also observed low (although statistically significant) correlations in expression fold changes between pairs of species, a result that was independent of the ability to correctly classify genes as differentially expressed. These results were based on conserved, high confidence ortholog groups that excluded complex families with duplication events shared by two or more species. Since many of the previously described *H. vulgare* cold responsive genes belonged to such complex families, we specifically investigated the regulation of these previously described genes using all ortholog groups, and again found that few genes displayed shared cold response across all species (Fig. 3). Taken together, our findings appear consistent with the lineage specific hypothesis of Pooideae cold adaptation, where cold induced regulation to a substantial degree has been gained or lost independently in the different lineages. However, it is unclear to what extent the most recent common ancestor (MRCA) of the Pooideae possessed response to cold, since we cannot determine whether lineage specific gain of cold response was more (or less) prominent than lineage specific loss.

The drastic cold stress during the E-O transition was likely an important cause for the evolution of cold adaptation in Pooideae. Previous studies have shown that many temperate plant lineages emerged during the E-O transition [25] and that the expansion of well-known cold responsive gene families in Pooideae coincided with this transition [15, 26]. From the dated phylogeny (Fig. 1a) as well as from earlier studies of the Pooideae phylogeny [2, 28], it is clear that all major Pooidaee lineages, including the core Pooideae, had emerged by the late Eocene. Hence, the five lineages studied here experienced the E-O transition as individual lineages (Fig. 1c). The observation that the five Pooideae lineages emerged during a relatively warm period before the E-O transition, that these species harbored high numbers of cold responsive genes specific to only one or two species and that the shared cold response genes showed no phylogenetic pattern, together suggests that a significant portion of the cold response in Pooideae lineages were gained during the last 40 M years. During this period, temperatures were constantly decreasing and dramatic cooling events took place, such as the E-O transition and the current Quaternary Ice Age. This must have placed a very strong selection pressure on plants favoring the evolution of cold adaptations, and would have acted on already diverged lineages and not on any ancestor of these lineages. However, although the independent gain of cold response would imply that the ancestor of the Pooideae possessed limited molecular cold response, further studies involving more species would be needed to definitively conclude on this issue.

Our results suggest that the Pooideae lineages evolved cold response programs that included a significant fraction of non-orthologous genes. This conclusion was reached both based on the conserved, high confidence ortholog groups, and based on a set of previously identified cold response genes (Fig. 3), and implies that these genomes contain many functionally redundant genes that can be co-opted in different combinations into the functional cold response program of the Pooideae. Gain and loss of protein coding genes might also have played an important role in the evolution of cold response of these species, however, here we focused on the gain and loss of regulatory response to cold in genes that were widely conserved across the phylogeny. It is worth noticing that although we observed many species specific cold response genes, all species pairs displayed a statistically significant correlation in cold response across all HCOGs (Fig. 2b) and a statistically significantly overlap in cold responsive genes (Fig. 2c). This could reflect that some genes are associated with biochemical functions more important or more suited for cold response than other genes [39, 40], and that different species thus have ended up co-opting orthologous genes into their cold response program more often than expected by chance.

### An adaptive potential in the Pooideae ancestor

Multiple independent origins of cold adaptation raise the question whether connecting traits exists in the evolutionary history of the Pooideae that can explain why the Pooideae lineages were able to shift to the temperate biome. The genes that showed a conserved cold response across all five species (Table 1) might have gained cold responsiveness in the MRCA. Subsequently, these genes might have increased the potential of Pooideae lineages to adapt to a cold temperate climate. Nine of these conserved cold response genes are involved in response to abiotic stresses in other plants, such as osmotic stress and drought. The *SNAC1-like / NAC transcription factor 67* is one example, with homologs induced by osmotic stress in the three monocts *Musa* × *paradisiaca* [41, 42], *Oryza sativa* [43] and *Triticum aestivum* [44]. Co-option of such genes into a cold-responsive pathway might have been the key to acquire cold tolerance. In fact, other studies have implied that drought tolerance might have facilitated the shift to temperate biomes [26, 45, 46]. Interestingly, most of the conserved genes were short-term cold responsive (Table 1) and this observation strengthens the hypothesis that existing stress genes might have been the first to be co-opted into the cold response program.

Also, three of the conserved cold responsive genes (GIGANTEA, PRR95 and AtPRR3-like) were associated with the circadian clock that is known to be affected by cold [47–49]. This might suggest that clock genes have had an important function in the Pooideae cold adaptation, for example by acting as a signal for initiating the cold defense. More generally, transcripts involved in photosynthesis and response to abiotic stimuli were significantly enriched among the genes with cold response in all species (Fig. 4a). An expanded stress responsiveness towards cold stress and the ability to down-regulate the photosynthetic machinery during cold temperatures to prevent photoinhibition might have existed in the early evolution of Pooideae. In conclusion, the conserved stress response genes discussed here may have represented a fitness advantage for the Pooideae ancestor in the newly emerging environment with incidents of mild frost, allowing time to evolve the more complex physiological adaptations required to endure the temperate climate with strong seasonality and cold winters that emerged following the E-O transition [23]. Consistent with this, Schubert et al. (unpublished) showed that the fructan synthesis and ice recrystallization inhibition protein gene families known to be involved in cold acclimation in core Pooideae species [10] evolved around the E-O split, whereas also earlier evolving Pooideae species show capacity to cold acclimate.

### Evolution of coding and regulatory sequences

The molecular mechanisms behind adaptive evolution are still poorly understood, although it is now indisputably established that novel gene regulation plays a crucial role [50]. The evolution of gene regulation proceeds by altering non-coding regulatory sequences in the genome, such as (cis-) regulatory elements [51], and has the potential to evolve faster than protein sequence and function. The high number of species specific cold response genes observed in this study is thus most consistent with the recruitment of genes with existing cold tolerance functions by means of regulatory evolution. However, previous studies have also pointed to the evolution of coding sequences [27] as underlying the acquisition of cold tolerance in Pooideae. To investigate possible coding evolution, we tested for the enrichment of positive selection among branch specific cold responsive genes (Fig. 4b). Although not statistically significant, there was a tendency for positive selection in genes gaining cold response in a period of gradual cooling preceding the E-O event. Thus, we saw evidence of both coding and regulatory evolution playing a role in cold adaptation in Pooideae, and that these processes may have interacted. Finally, gene family expansion has previously been implied in cold adaptation in Pooideae [15, 26]. As previously discussed, the conservative filtering of ortholog groups employed in this study removed complex gene families containing duplication events shared by two or more species. Interestingly, out of the 33 previously described *H. vulgare* cold responsive genes (Fig. 3), as many as 22 were not included in the high confidence ortholog groups, the main reason being that they belonged to gene families with duplications involving two or more species. This observation thus confirms that duplication events are a relatively common feature of cold adaptation. Although *de novo* assembly of transcriptomes from short-read RNA-Seq data is a powerful tool that has vastly expanded the number of target species for conducting transcriptomic analysis, the approach has limited power to distinguish highly similar transcripts such as paralogs. Further insight into the role of duplication events in Pooideae cold adaptation would therefore benefit immensely from additional reference genomes.

## Conclusion

Here we investigated the cold response of five Pooideae species, ranging from early diverging lineages to core Pooideae species, to elucidate evolution of adaptation to cold temperate regions. We observed extensive lineage specific cold response that seems to have evolved chiefly after the *B. distachyon* lineage and the core Pooideae diverged, possible initiated by the drastic temperature drop during the E-O transition. However, we do also see signs of conserved response that potentially represents a shared potential for cold adaptation that explain the success of Pooideae in temperate regions. This included several general stress genes with conserved short-term response to cold as well as the conserved ability to down-regulate the photosynthetic machinery during cold temperatures. Taken together, our observations are consistent with a scenario where the biochemical functions needed for cold response were present in the Pooideae common ancestor, and where different Pooideae lineages have assembled, in parallel, different overlapping subsets of these genes into fully functional cold response programs through the relatively rapid process of regulatory evolution. Answering whether gain or loss of cold response was most prominent in the evolution of cold adaption in these species, and thus whether the last common ancestor of the Pooideae possessed extensive cold response or not, will require further studies.

## Methods

### Plant material, sampling and sequencing

To address our hypothesis, we selected five species to cover the phylogenetic spread of Pooideae. The selected species also represent major, species rich lineages or clades in the Pooideae subfamily, or belong to very early diverging lineages [5]. Seeds were collected either in nature or acquired from germplasm collections for the following five species:

*Nardus stricta*, (2n=2x=26) is a perennial species, distributed in Europe, western parts of Asia, and North Africa and introduced to New Zealand and North America. Seeds were collected in July 2012 in Romania, [46.69098, 22.58302].

*Melica nutans* L. (2n=2x=18) is a perennial species distributed in temperate parts of Eurasia [52– 54]. Seeds were collected in Germany, [50.70708, 11.23838], in June 2012.

S*tipa lagascae* Roem & Schult. (2n=2x=22) is a perennial species that is distributed in temperate regions around the Mediterranean Sea and parts of temperate West Asia. Seeds from accession PI 250751 were acquired from U.S. National Plant Germplasm System (U.S.-NPGS) via Germplasm Resources Information Network (GRIN),

*Brachypodium distachyon* (L.) P. Beauv. (2n=2x=10) is an annual species natively distributed in Europe, East Africa and temperate parts of West Asia. Seeds for accession ‘Bd1-1’ (W6 46201) were acquired from U.S.-NPGS via GRIN.

*Hordeum vulgare* L. (2n=2x=14) seeds for cultivar ‘Igri’ were provided by Prof. Åsmund Bjørnstad, Department of Plant Sciences, Norwegian University of Life Sciences, Norway.

Seeds were germinated and initially grown in a greenhouse at a neutral day length (12 hours of light), 17°C and a minimum artificial light intensity of 150 μmol/m2s. To ensure that individual plants were at comparable developmental stages at sampling, the onset of treatment for different species were based on developmental stages rather than absolute time. Most importantly, none of the plants had transitioned from vegetative to generative phase, as this could have affected the cold response. Plants were grown until three to four leaves had emerged for *M. nutans*, *S. lagascae*, *B. distachyon* and *H. vulgare*, or six to seven leaves for *N. stricta* (which is a cushion forming grass that produces many small leaves compared to its overall plant size). Depending on the species, this process took one (*H. vulgare*), three (*B. distachyon* and *S. lagascae*), six (*M. nutans*) or eight (*N. stricta*) weeks from the time of sowing. Subsequently, plants from all species were randomized and distributed to two cold chambers with short day (8 hours of light), constantly 6°C and a light intensity of 50 μmol/m2s. Plants were kept in cold treatment for the duration of the experiment. Leaf material for RNA isolation was collected i) in the afternoon (at zeitgeber (ZT) 8) on the day before cold treatment (D0) and in the afternoon (ZT 8) on the first day after cold treatment (after 8 hours of cold treatment, D1) and ii) in the morning (ZT 0) before cold treatment (W0), 4 weeks after cold treatment (W4) and 9 weeks after cold treatment (W9) (Fig. 1d). The sampling time points were chosen to be able to separate chilling stress responses (first day of treatment) and long-term responses that represent acclimation to freezing temperatures (4 and 9 weeks). Flash frozen leaves were individually homogenized using a TissueLyser (Qiagen Retsch) and total RNA was isolated (from each leaf) using RNeasy Plant Mini Kit (Qiagen) following the manufacturer’s instructions. The purity and integrity of total RNA extracts was determined using a NanoDrop 8000 UV-Vis Spectrophotometer (Thermo Scientific) and 2100 Bioanalyzer (Agilent), respectively. For each time point, RNA extracts from five leaves sampled from five different plants were pooled and sequenced as a single sample. In addition, replicates from single individual leaves were sequenced for selected timepoints (see Table S1 and “Differential expression” below). Two time points lacked expression values: W9 in B. distachyon (RNA integrity was insufficient for RNA sequencing) and W0 in S. lagascae (insufficient supply of plant material). Samples were sent to the Norwegian Sequencing Centre, where strand-specific cDNA libraries were prepared and sequenced (paired-end) on an Illumina HiSeq 2000 system. The raw reads are available in the ArrayExpress database (www.ebi.ac.uk/arrayexpress) under accession number E-MTAB-5300.

### Transcriptome assembly and ortholog inference

Using Trimmomatic v0.32 [55], all reads were trimmed to a length of 120 bp, Illumina TruSeq adapters were removed from the raw reads, low quality bases were trimmed using a sliding window of 40 bp and an average quality cut-off of 15 and reads below a minimum length of 36 bp were discarded. Read quality was checked using fastqc v0.11.2. For each species, transcripts were assembled *de novo* with Trinity v2.0.6 [56] (strand specific option, otherwise default parameters) using reads from all samples. Coding sequences (CDS) were identified using TransDecoder rel16JAN2014 [57]. Where Trinity reported multiple isoforms, only the longest CDS was retained. Ortholog groups (OGs) were constructed from the five *de novo* transcriptomes and public reference transcriptomes of *H. vulgare* (barley_HighConf_genes_MIPS_23Mar12), *B. distachyon* (brachypodium v1.2), *O. sativa* (rap2), *Z. mays* (ZmB73_5a_WGS)*, S. bicolor* (sorghum 1.4) and *L. perenne* (GenBank TSA accession GAYX01000000) using OrthoMCL v2.0.9 [58]. All reference sequences except *L. perenne* were downloaded from http://pgsb.helmholtz-muenchen.de/plant/plantsdb.jsp. A summary of the results is provided in Table S1.

### High confidence ortholog groups

To compare gene expression across Pooideae, we required ortholog groups containing one gene from each species that all descended from a single gene in the Pooideae ancestor. As the ortholog groups (OGs) inferred using orthoMCL sometimes cluster more distantly related homologs as well as include both paraphyletic and monophyletic paralogs, we further refined the OGs by phylogenetic analysis. Several approaches to phylogenetic refinement has been proposed previously (see e.g. [59]). Here we first aligned protein sequences within each OG using mafft v7.130 [60] and converted to codon alignments using pal2nal v14 [61]. Gene trees were then constructed from the codon alignments using Phangorn v1.99.14 [62] (maximum likelihood GTR+I+G). Trees with apparent duplication events before the most recent common ancestor of the included species were split into several trees. This was accomplished by identifying in-group (Pooideae) and out-group (*Z. mays*, *S. bicolor* and *O. sativa*) clades in each tree, and then splitting the trees so that each resulting sub-tree contained a single out-group and a single in-group clade. Finally, we only retained the trees were all species in the tree formed one clade each (i.e. only monophyletic paralogs), *B. distachyon* and *H. vulgare* formed a clade and at least three of the five studied species were included. These trees constituted the high confidence ortholog groups (HCOGs).

*De novo* assembly followed by ortholog detection resulted in higher numbers of monophyletic species-specific paralogs than the number of paralogs in the reference genomes of *H. vulgare* and *B. distachyon*. This apparent overestimation of paralogs was most likely the result of the *de novo* procedure assembling alleles or alternative transcript isoforms into separate contigs. We also observed some cases where the number of paralogs were under-estimated compared to the references, which may be due to low expression of these paralogs or the assembler collapsing paralogs into single contigs. Since the *de novo* assembly procedure did not reliably assemble paralogs, we chose to represent each species in each HCOG by a single read-count value equal to the sum of the expression of all assembled paralogs. By additionally setting counts for missing orthologs to zero, we created a single cross species expression matrix with HCOGs as rows and samples as columns (Table S3).

### Species tree

Ortholog groups with a single ortholog from each of the five *de novo* Pooideae species and *O. sativa* (after splitting the trees, see “High confidence ortholog groups”) were used to infer dated gene trees. To this end, BEAST v1.7.5 [63] was run with an HKY + Γ nucleotide substitution model using an uncorrelated lognormal relaxed clock model. A Yule process (birth only) was used as prior for the tree and the monophyly of the Pooideae was constrained. Prior estimates for the *Oryza*-Pooideae (53 Mya [SD 3.6 My], [37]) and *Brachypodium*-*Hordeum* (44.4 Mya [SD 3.53 My], [28]) divergence times were used to define normally distributed age priors for the respective nodes in the topology. MCMC analyses were run for 10 million generations and parameters were sampled every 10.000 generation. For each gene tree analysis, the first 10 percent of the estimated trees were discarded and the remaining trees were summarized to a maximum clade credibility (MCC) tree using TreeAnnotator v1.7.5. The topology of the species tree was equal to the most common topology among the 3914 MCC trees, with internal node ages set equal to the mean of the corresponding node age distributions of the MCC gene trees.

### Differential expression

Reads were mapped to the *de novo* transcriptomes using bowtie v1.1.2 [64], and read counts were calculated with RSEM v1.2.9 [65]. In HCOGs, read counts of paralogs were summed (analogous to so called monophyly masking [66]) and missing orthologs were assumed to not be expressed (i.e. read counts equal to zero). To identify conserved and diverged cold response across species, we probed each HCOG for differentially expressed genes (DEGs). Specifically, DEGs were identified using DESeq2 v1.6.3 [67] with a model that combined the species factor and the timepoint factor (with timepoints W4/9 as a single level). Pooled samples provided robust estimates of the mean expression in each time point. To also obtain robust estimates of the variance, the model assumed common variance across all timepoints and species within each HCOG, thus taking advantage of both biological replicates available for individual time points within species and the replication provided by analysing several species. For each species, we tested the expression difference between D0 and D1 (short-term response) and the difference between W0 and W4/9 (long-term response) (Fig. 1d). *B. distachyon* lacked the W9 samples and long-term response was therefore based on W4 only. *S. lagascae* lacked the W0 sample and long-term response was therefore calculated based on D0. As a result, the observed diurnal effect (Fig. 2a) might have resulted in more unreliable estimates of the long term cold response in *S. lagascae* since for this species the afternoon sample (D0) was used to replace the missing morning sample (W0). Genes with a false discovery rate (FDR) adjusted p-value < 0.05 and a fold change > 2 were classified as differentially expressed.

### Sample clustering

Sample clustering was based on read counts normalized using the variance-stabilizing transformation (VST) implemented in DESeq2 (these VST-values are essentially log transformed). HCOGs that lacked orthologs from any of the five species, or that contained orthologs with low expression (VST < 3), were removed, resulting in 4981 HCOGs used for the clustering. To highlight the effect of the cold treatment over the effect of expression level differences between species, the expression values were normalised per gene and species: First, one expression value was obtained per timepoint per gene by taking the mean of the replicates. Then, these expression values were centered by subtracting the mean expression of all timepoint. Distances between all pairs of samples were calculated as the sum of absolute expression difference between orthologs in the 4981 HCOGs (i.e. manhattan distance). The tree was generated using neighbor-joining [68].

### Comparison with known cold responsive genes

A set of *H. vulgare* genes independently identified as cold responsive were acquired from supplementary table S10 in [38]. These genes were found to be responsive to cold in three independent experiments with Plexdb accessions BB64 [69], BB81 (no publication) and BB94 [38]. The probesets of the Affymetrix Barley1 GeneChip microarray used in these studies were blasted (blastx) against all protein sequences in our OGs. Each probe was assigned to the OG with the best match in the *H. vulgare* reference. If several probes were assigned to the same OG, only the probe with the best hit was retained. Correspondingly, if a probe matched several paralogs within the same OG, only the best match was retained. DESeq2 was used to identify short-term response DEGs for all transcripts in all OGs (i.e. this analysis was not restricted to the HCOGs), and these were compared to DEGs from [38]. The statistical significance of the overlap between our results and those reported in [38] was assessed for each species by counting the number of genes that had the same response (up-or down-regulated DEGs) and comparing that to a null distribution. The null distribution was obtained from equivalent counts obtained from 100 000 trials where genes were randomly selected from all expressed genes (mean read count > 10) with an ortholog in *H. vulgare*.

### Gene ontology enrichment tests

Gene Ontology (GO) annotations for *B. distachyon* were downloaded from Ensembl Plants Biomart and assigned to the HCOGs. The TopGO v2.18.0 R package [70] was used to calculate statistically significant enrichments (Fisher’s exact test, p < 0.05) of GO biological process annotations restricted to GO plant slim in each set of branch specific DEGs using all annotated HCOGs as the background. Branch specific DEGs were those genes that were exclusively differentially expressed in all species within a clade in the phylogenetic tree.

### Positive selection tests

Each of the HCOGs were tested for positive selection using the branch-site model in codeml, which is part of PAML v4.7 [71]. We only tested branches for positive selection in HCOGs meeting the following criteria: (i) The tested branch had to be an internal branch also in the gene tree (i.e. there was at least two species below the branch). (ii) The species below and above the tested branch in the gene tree had to be the same as in the species tree or a subset thereof. (iii) The first species to split off under the branch had to be the same as in the species tree (for the early split, either *S. lagascae* or *M. nutans* was allowed). We then used the Hypergeometric test to identify statistically significant overrepresentation of positive selection among branch specific DEGs (see “Gene ontology enrichment tests”) at the Pooideae base (PB), the early split (ES) and the late split (LS) branches.

## Declarations

### Availability of data and material

The raw reads, the assembled transcripts and the raw read counts are available in the ArrayExpress database (www.ebi.ac.uk/arrayexpress) under accession number E-MTAB-5300.

### Authors’ contributions

All authors designed the experiment. M.S. performed the growth experiments, sampled and prepared RNA for sequencing, helped designing the data analysis pipeline, contributed to the positive selection analysis and performed the phylogenetic analyses. L.G. developed, implemented and conducted the data analysis. All authors interpreted the results. M.S. and L.G. wrote the manuscript with input from S.F., T.R.H., and S.R.S.

## Acknowledgements

We thank Åsmund Bjørnstad and USDA-NPGS GRIN for providing seeds of *H. vulgare*, and *B. distachyon* and *S. lagascae*, respectively. For technical assistance handling plants during growth experiments we thank Øyvind Jørgensen. We are grateful to Erica Leder, Thomas Marcussen, Ursula Brandes and Camilla Lorange Lindberg for helpful comments on earlier versions of this manuscript.

## Supporting Information

**Table S1. Summary statistics for the sampling, the transcriptome assembly, coding sequence detection and ortholog group inference**. Isoforms were not included in the counts. Only ortholog groups with at least one coding transcript from the five studied species were included.

**Table S2. High confidence ortholog groups (HCOGs**). HCOGs generated by filtering and splitting ortholog groups (see Methods). Ortholog groups were stored as a table with the ortholog group IDs as rows and species as columns. Each cell contains sequence IDs separated by “,”. Groups that were the result of splitting larger ortholog groups were marked by a number suffix in the group ID.

**Table S3. A cross species expression matrix.** Combined read counts for high confidence ortholog groups. Column represents samples and rows represents HCOGs. The sample IDs in the column header consists of the species ID, the time point, indication of whether the sample is pooled from five individual plants (“mix”) or just a single individual plant (“ind”) and the replicate number.

**Table S4. Differential expression results for the high confidence ortholog groups (HCOGs).** A table with rows representing HCOGs and columns representing differential expression results including log2 fold changes, P-values and FDR adjusted p-values for short-and long-term responses.

**Figure S1: The four most common gene tree topologies.** We generated gene trees from a selected set of 3914 ortholog groups (see Methods). This figure depicts the four most common topologies.

